# *In vitro* hepatitis C virus infection and hepatic choline metabolism

**DOI:** 10.1101/746776

**Authors:** Kaelan Gobeil Odai, Conor O’Dwyer, Rineke Steenbergen, Tyler A. Shaw, Tyler M. Renner, Peyman Ghorbani, Mojgan Rezaaifar, Shauna Han, Marc-André Langlois, Angela M. Crawley, Rodney S. Russell, John P. Pezacki, D. Lorne Tyrrell, Morgan D. Fullerton

**Affiliations:** Department of Biochemistry, Microbiology and Immunology, Faculty of Medicine, University of Ottawa, Ottawa, Canada; University of Ottawa Centre for Infection, Immunity and Inflammation and Centre for Catalysis Research and Innovation; Li Ka Shing Institute of Virology, Department of Medical Microbiology and Immunology, University of Alberta, Edmonton, Canada; Department of Chemistry and Biomolecular Sciences, Faculty of Science, University of Ottawa, Ottawa, Canada; Chronic Disease Program, Ottawa Hospital Research Institute, Ottawa, Canada; Department of Medicine, Division of Infectious Diseases, The Ottawa Hospital, Ottawa, Canada; Department of Biology, Faculty of Science, Carleton University, Ottawa, Canada; Immunology and Infectious Diseases, Faculty of Medicine, Memorial University of Newfoundland, St. John’s, Canada

**Author notes:** Correspondence (M.D.F.).

**Keywords:** choline, Hepatitis C Virus (HCV), phosphatidylcholine, virus, Huh7.5, CTL1, SLC44A1, immunometabolism

## Abstract

Choline is an essential nutrient required for normal neuronal and muscular development, as well as homeostatic regulation of hepatic metabolism. In the liver, choline is incorporated into the main eukaryotic phospholipid, phosphatidylcholine (PC) and can enter one carbon metabolism via mitochondrial oxidation. Hepatitis C virus (HCV) is a hepatotropic positive-strand RNA virus that similar to other positive-strand RNA viruses can impact phospholipid metabolism. In the current study we sought to interrogate the link between choline transport and early HCV infection. Namely, we aimed to investigate how HCV modulates markers of choline metabolism following *in vitro* infection, while subsequently assessing how the inhibition of choline uptake and metabolism upon concurrent HCV infection may alter early viral replication. Finally, we assessed whether these parameters were consistent between cells cultured in fetal bovine serum (FBS) or human serum (HS), conditions known to differentially affect *in vitro* HCV infection. We observed that choline transport in FBS-cultured Huh7.5 cells is facilitated by the intermediate affinity transporter choline transporter-like family (CTL), and that CTL1 expression and the incorporation of choline into PC is diminished in 24 h infected FBS-cultured cells. Reciprocally, limiting the availability of choline for PC synthesis resulted in increased HCV replication at this early stage. In chronically HS-cultured Huh7.5 cells, there were no differences in the expression of choline transporters upon HCV infection or alterations to viral replication when choline transport was inhibited compared to control treatments. However, inhibiting choline uptake and metabolism in this system significantly impaired the production of infectious virions in HS-cultured cells. These results suggest that in addition to a known role of choline kinase, the transport of choline, potentially via CTL1, might also represent an important and regulated process during HCV infection.

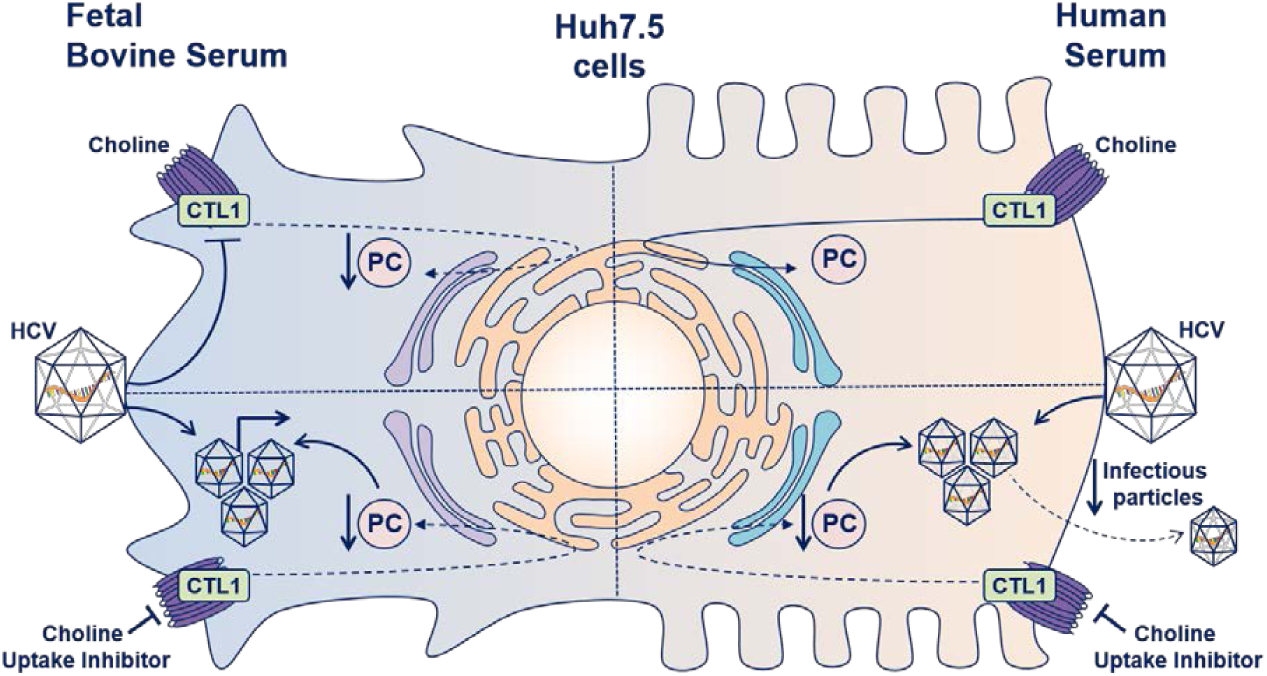

## 1 Introduction

Hepatitis C virus (HCV) is a positive-strand (+)RNA virus that is inherently primate-borne and principally targets the liver [1]. HCV is part of the *Flaviviridae* family along with other viruses such as Yellow fever virus, West Nile virus, Dengue virus and Zika virus [2]. Approximately 71 million people worldwide are infected with HCV, or approximately 1% of the global population [3]. Although some cases only lead to mild illness, most infected individuals (75-85%) develop chronic HCV infection. While recent advancements in direct-acting antiviral therapy have proven to be highly efficacious (∼ 98% cure rates), barriers to therapy access, viral resistance and low diagnosis rates minimizing efforts towards virus elimination, chronic HCV infection remains a significant medical concern. With no widely available vaccine, and culminating in chronic hepatitis, cirrhosis, as well as one of the world’s leading causes of death, hepatocellular carcinoma (HCC) [4], HCV infection has important implications in global health outcomes. Together with non-alcoholic fatty liver disease, Hepatitis C infection are the most common indications for a liver transplant in the Western world [3].

Crucial throughout all stages of development, choline plays an essential role in many, if not all tissues [5-7]. While choline is specifically taken into neuronal tissues via the high-affinity choline transporter (CHT1/*SLC5A7*), in non-neuronal tissues, low- and intermediate-affinity choline transport systems have been identified [8]. Organic cation transporters (OCT1-3/*SLC22A1-3*) represent a family of low-affinity and promiscuous cation transporters. Intermediate-affinity transport is mediated by the choline transporter-like protein family (CTL1-5/*SLC44A1-5*), where CTL1 has been implicated as the main choline transporter in most tissues [9].

Phosphatidylcholine (PC) synthesis is the primary fate of hepatic choline, which accounts for approximately half of the phospholipid species in most mammalian cells [10]. Although in the liver and kidney, choline serves as a methyl group donor via its mitochondrial oxidation to betaine [11]. In non-hepatic tissues, PC is exclusively made through the well-described cytidine diphosphate (CDP)-choline (or Kennedy) pathway; however, in the liver, PC supply is supplemented by the phosphatidylethanolamine (PE) *N*-methyl transferase (PEMT) pathway [12-14]. This pathway is thought to supply up to 30% of hepatic PC by methylating and converting PE to PC; however, it cannot fully compensate for disruptions in the CDP-choline pathway [15].

Previous studies have demonstrated that (+)RNA viruses impact cellular phospholipid metabolism [16,17], and certain viruses have been observed to increase the PC content in distinct cellular membranes during sustained infection [18]. Interestingly, the role of choline kinase alpha (CHKα), the first step of the CDP-choline pathway, has proven to be a requisite component in maintaining the integrity of the HCV membranous web, the site of viral replication located on the ER [19,20]. Moreover, CHKα was shown to be essential in shuttling the CHKα-PIKIIIα-NS5A ternary HCV assembly complex to the viral replication complex, giving further credence that PC metabolism may be integral to the HCV life cycle [21]. Finally, PC forms the principle membrane component of hepatocyte-derived ApoB-containing lipoproteins such as very low-density lipoproteins (VLDL). It has been previously shown that HCV subverts VLDL to make lipoviral particles that subsequently undergo a multi-step endocytosis that aides in HCV infection of hepatocytes [22-25]. Interestingly, it has recently been shown that Huh7.5 cells, which represent one of the most well-characterized cell culture models for studying hepatic viral biology, behave very differently when cultured in human serum (HS) compared to normal fetal bovine serum (FBS) [25,26]. HS-cultured Huh7.5 cells are more polarized, express more functional hepatocyte markers and dramatically increase VLDL secretion. Although some aspects of choline metabolism have been investigated in sustained infection models, the importance of choline transport and the immediate cellular response of PC metabolism to the initial HCV infection have yet to be described.

Therefore, in this study we sought to interrogate this potential link and observed that the CTL family of transporters mediated the functional uptake of choline in Huh7.5 cells. Concordant with decreases in choline uptake and CTL1 expression, the rate of flux through the CDP-choline pathway was lower after 24 h of HCV infection in FBS-cultured cells. The inhibition of choline transport during early infection in FBS-cultured cells augments viral replication and finally, HS-cultured cells do not directly phenocopy FBS-cultured conditions. Our results suggest that choline transport represents a previously underappreciated aspect of PC metabolism that is both able to regulate and be regulated by HCV infection in FBS- and HS-cultured Huh7.5 cells.

## 2 Materials and Methods

### 2.1. Huh7.5 cell culture

Huh7.5 cells were a kind gift from C.M. Rice (Rockefeller University). FBS-cultured cells were cultured in Dulbecco’s Modified Eagle Media (DMEM) containing 10% FBS (Wisent) and 1% penicillin/streptomycin (Gibco). Cells were maintained under 100% confluency and were split using standard trypsin/EDTA protocol. Cells of different passages on different days were considered as biological replicates. HS-cultured cells were grown as previously described [25], where 2% HS (Valley Biomedical) was used in place of FBS. These cells form confluent layers of growth arrested cells, that do not need further sub-culturing.

### 2.2. JFH-1 HCV infection protocol

The initial harvest of JFH-1 HCV was following electroporation into FBS-cultured cells, as previously described [27]. Viral production (RNA/mL and 50% tissue culture infectious dose [TCID50]/mL) was further monitored. Supernatants were collected after 4 days and these viral stocks were either used to propagate more FBS-cultured virus or for HS-cultured infection experiments described below. For FBS-cultured cells, Huh7.5 cells were plated at 70% confluency and infected with live JFH-1 strain HCV virions at a MOI = 1 for 4 h containing 5% lipoprotein deficient serum. Fully differentiated and confluent HS cultured cells were infected at an MOI = 1. Following this 4 h incubation, the media was removed and cells were washed 3 times before being cultured in FBS or HS media. Experiments were performed 24 h after infection for FBS cells and 92 hours after infection for HS cells.

### 2.3. JFH-1 HCV titration

Infectious virus particles were quantified using a focus forming assay. Virus supernatant was serially diluted 10-fold in serum-free medium and dilutions were used to infect Huh7.5 cells seeded (at 5×10^4^ per well) onto 8-well Lab-Tek II chamber slides (NUNC) for 4 h. Following incubation, the infectious medium was removed and replaced with fresh medium containing 10% FBS and 1X NEAA. Forty-eight hours post infection, cells were washed with 1X PBS and fixed with acetone for 5 minutes before staining with HCV core monoclonal antibody (1:100; Thermo Fisher Scientific; MA1080), followed by a secondary antibody, Alexa Fluor 488– conjugated goat anti-mouse (1:250; Thermo Fisher Scientific; A-11029). Viral titers are expressed as the number of focus-forming units (FFU) per mL of supernatant.

### 2.4. [^3^H]-Choline uptake experiments

The uptake of choline in Huh7.5 cells was determined by using [^3^H]-choline (Perkin Elmer) to evaluate uptake the desired time points. Prior to the addition of radiolabeled choline, Huh7.5 cells seeded in 24-well plates were washed twice with PBS then incubated with Krebs-Ringer-HEPES buffer (KRH; 130 mM NaCl, 1.3 mM KCl, 2.2 mM CaCl_2_, 1.2 mM MgSO_4_, 1.2 mM KH_2_PO_4_, 10 mM HEPES pH 7.4, 10 mM glucose) for one h at 37°C prior to treatment to remove extracellular choline. Following KRH incubation, [^3^H]-choline (1 µCi/mL in KRH) was added to the cells for the desired time points (0-30 minutes) and incubated at 37°C. For uptake kinetics, cells were incubated in increasing concentrations of non-radiolabeled choline for 10 min. The [^3^H]-choline was then removed, and the cells were washed twice with KRH buffer, then lysed in 300 µL of 0.1 M NaOH. Cells were flash frozen using liquid nitrogen, thawed then the lysate was scraped and collected. Cell lysate was centrifuged at 20,000 × g for 5 min and supernatant was collected. Cellular protein amounts were determined by BCA protein quantification assay according to manufacturer’s instructions (Thermo Fisher Scientific) and radioactivity was quantified through liquid scintillation counting (LSC). Choline uptake and kinetics were calculated as previously described [28].

### 2.5. [^3^H]-Choline incorporation into PC

Huh7.5 cells were seeded in 12-well plates and incubated at 37°C up to 8 h with DMEM containing 0.5 µCi/mL of [^3^H]-choline. Cells were then washed twice with PBS and 250 µL of PBS was the added to the cells, which were then flash frozen in liquid nitrogen. Following the collection of cell lysate, of which a 50 µL aliquot was counted by LSC, we performed a total lipid extraction protocol described by Bligh and Dyer [29]. The organic phase was evaporated under nitrogen and resuspended in 25 µL of chloroform to concentrate the lipid species. The concentrated solution was then subjected to thin layer chromatography as previously described and radioactivity corresponding to PC was counted by LSC [30].

### 2.6. Choline inhibition by HC3

FBS-cultured Huh7.5 cells were infected as described above in the presence or absence of 20 or 200 μM HC3 for 24 h. Following treatment, cells were either processed for RNA or labeled with [^3^H]-choline as described above. HS-cultured Huh7.5 cells were infected as described above and treated with 20 or 200 μM HC3 at 72 h post-infection, for a total of 24 h before being harvested for RNA quantification.

### 2.7. RNA isolation and quantification

Total RNA was extracted using the TriPure reagent protocol (Roche Life Sciences). Following extraction, the RNA was resuspended in 20 µL of RNAse/DNAse-free H_2_O (Wisent) and concentration equalized. RNA was then reversed transcribed using the QuantiNova™ kit (Qiagen) according to the kit’s instructions. Transcript expression was determined by using primers obtained from PrimerBank, designed through NCBI PrimerBlast, or from previous published studies [31] (Table 1). These primers were used in conjunction with the BrightGreen 2x qPCR mastermix (ABM). The relative transcript expression was determined using the delta-delta *Ct* method [32] and normalized to the averages of *β-actin, HSP-90*, and *β*_*2*_*-microglobulin*. The efficiencies of all primers were validated before use. For quantification of viral RNA, infected cells underwent total RNA extraction as described above. Reverse transcription was initiated using the OneScript cDNA Synthesis Kit (ABM) as per the kit instructions, using random hexamer primers. Viral RNA was assess using previously published primers (Table 1) and shown relative to *HSP-90* and *β actin*. TCID50s were determined as described [33].

### 2.8. Immunoblotting

To investigate protein expression, cells were lysed in native lysis buffer (50 mM Tris-HCl pH 7.4, 150 mM NaCl 1 mM EDTA, 100 μM sodium orthovanadate, and protease inhibitor cocktail tablet; Roche). Lysate was equalized and loaded onto an 8% denaturing SDS-PAGE gel. Native protein lysate samples were subsequently probed for CTL1 as well as β actin as a loading control. Following electrophoresis, gels were transferred using the Trans-Blot^®^ system (BioRad) onto PVDF membranes (1.3A for 17 min) using Bjerrum Schafer-Nielsen buffer (48mM Tris, 39mM glycine, 20% methanol). The membranes were blocked in 5% BSA for one h then incubated overnight at 4°C with primary rabbit antibody targeted towards CTL1 (Abcam #177877) or β actin. Following overnight incubation, the membranes were washed 4 times with TBS-T (20 mM Tris, 150 mM NaCl, 0.05% Tween^®^ 20) then incubated for an h with a 1:1000 dilution of HRP-conjugated anti-rabbit secondary antibody made in 5% BSA. The membranes were then again washed 4 times with TBS-T then treated with Clarity™ Western ECL solution (BioRad) according to manufacturer instructions. Treated membranes were visualized and imaged using LAS 4010 ImageQuant imaging system (General Electric).

### 2.9. Statistics

All statistical analyses were performed using Prism7 (GraphPad Software Inc.). Experiments consisting of only two groups were assessed by Student’s *t* test. Choline uptake kinetic measurements were fit to Michaelis-Menten equations and uptake inhibition curve was fit to a logarithmic(inhibitor) vs. response curve fit. Experiments involving two groups and multiple treatments were compared using two-way ANOVA with a Sidak *post-hoc* analysis. All data represent mean±SEM, unless specified in the figure legend.

## 3 Results

### 3.1. Determination of choline transport kinetics

To establish a kinetic profile of choline uptake in the Huh7.5 cell line cultured in FBS, we first performed uptake experiments using [^3^H]-choline and established that transport was linear over the course of 30 min (data not shown). Following this, we determined that the apparent affinity for choline (K_M_) was 66.8±9.08 µM and that the V_max_ was 296.3±14.45 µmol/min (Figure 1A), indicative of an intermediate affinity transport system [8,9]. There is evidence that hepatic choline transport is mediated by the CTL family, namely CTL1, as well as the low-affinity OCT family [8,9,34]. We next used hemicholinium-3 (HC3), a potent pharmacological inhibitor of high and intermediate affinity choline transporters, which revealed that ∼ 90% of choline transport was sensitive to HC3 uptake inhibition (Figure 1B). In addition to HC3, we also aimed to assess the potential contribution of the OCTs by using a well-characterized OCT inhibitor, quinine. At the highest dose (200 μM), choline uptake remained unchanged (Figure 1C).

**Figure 1.**
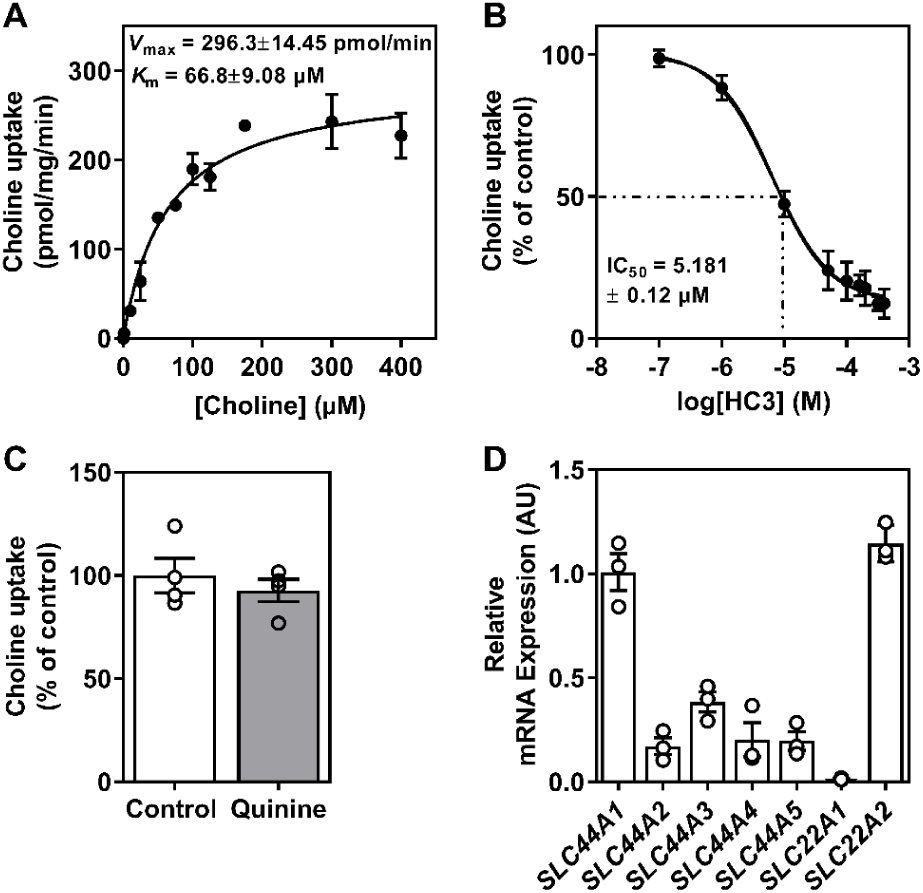
Characterization of Huh7.5 cell choline uptake. A) Choline uptake saturation kinetics fit to a Michaelis-Menten curve. B) Inhibition of choline uptake in response to hemicholinium-3 (HC3). C) Choline uptake inhibition in response to 200 μM quinine (OCT inhibitor). D) Relative transcript expression of choline transport genes, shown relative to *SLC44A1* and normalized to the average of *β actin, HSP90* and *β-microglobulin*. Data are mean±SEM and represent 3-4 independent experiments.

Finally, we profiled the transcript expression of the relevant choline transporters. The relative expression of CTL1 (SLC44A1) transcript was higher compared to that of other family members, while only OCT1 and OCT2 (SLC22A1 and SLC22A2) were detectable from the OCT family (Figure 1D). Additionally, to rule out the typically neuronal high affinity choline transporter CHT1/SLC5A7 as a contributor in FBS-cultured Huh7.5 cells, we confirmed that expression levels were undetectable (data not shown). Taken together, these results implicate an intermediate affinity transport system as the predominant choline transporters in FBS-cultured cells.

### 3.2. Effects of early HCV infection on choline transport and incorporation

To investigate the potential role of choline and PC metabolism in relation to HCV pathogenicity, we next aimed to determine how choline transport and metabolism would be affected by HCV infection in the early stages of *in vitro* infection. First, we compared non-infected, naïve FBS-cultured Huh7.5 cells, to those infected with the JFH-1 strain of HCV (MOI = 1). After 24 h, infected cells displayed a small but significant impairment in choline uptake and a 2-fold reduction in *V*_max_ (Figure 2A and 2B), with no change in apparent affinity for choline (*K*_M_). Accompanying this, 24 h HCV infection lowered [^3^H]-choline incorporation into PC compared to non-infected control cells (Figure 2C).

**Figure 2.**
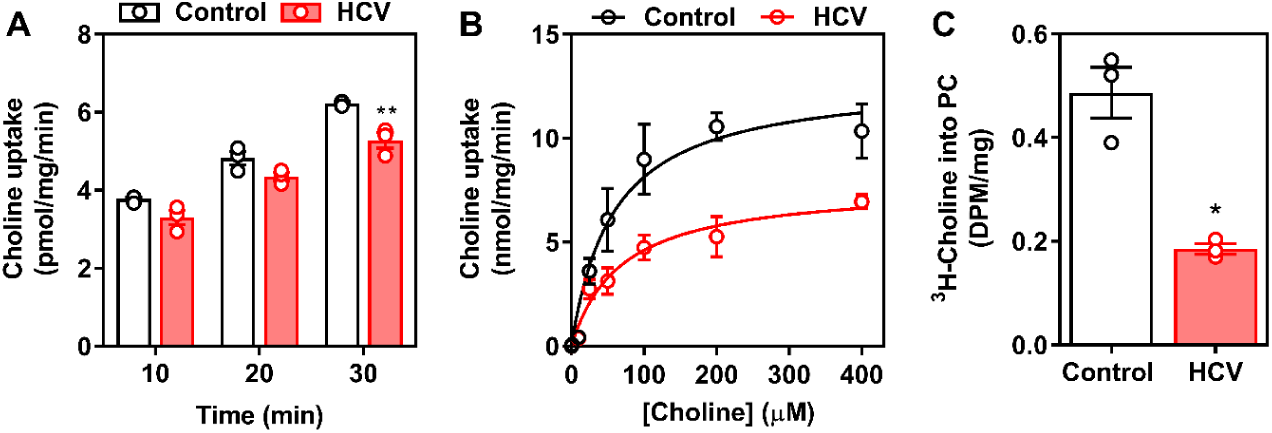
Choline uptake and incorporation 24 h post-infection. FBS-cultured Huh7.5 cells were infected at an MOI of 2 for 24 h. A) [^3^H]-choline uptake, B) choline uptake saturation kinetics fit to a Michaelis-Menten curve and C) [^3^H]-choline incorporation into PC was then determined. Data are mean±SEM and are representative of 3 independent experiments.

These measurements were associated with decreased transcript and protein expression of the choline transporter CTL1 (Figure 3A and 3B). However, there were no significant changes in the transcript expression of other choline transporters or CDP-choline pathway genes (Figure 3A).

**Figure 3.**
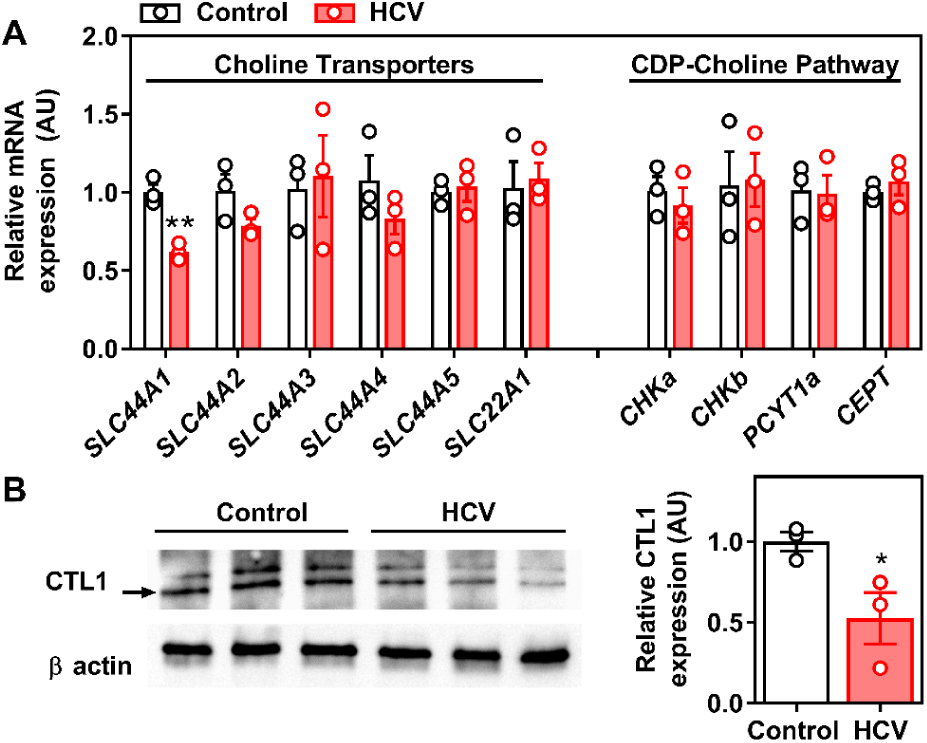
Choline transporter CTL1 expression at 24 h post-infection. FBS-cultured Huh7.5 cells were infected at an MOI of 2 for 24 h before cells were collected to measure A) choline transporter and CDP-choline pathway transcript expression, shown relative to uninfected control cells and normalized to the average of *β actin* and *HSP90* and B) CTL1 protein expression. Data are mean±SEM and represent 3 independent experiments.

### 3.3. Inhibiting choline uptake alters early HCV replication

We next aimed to ascertain if inhibiting choline availability would affect the early stages of HCV infection. Using HC3 as a pharmacological means of achieving choline transport inhibition, FBS-cultured Huh7.5 cells were infected with HCV as above (MOI = 1), but in the presence or absence of 20 or 200 μM HC3 for 24 h. As above (Figure 2C), in the absence of HC3, HCV infection resulted in a diminished incorporation of [^3^H]-choline into PC after 24 h. Treatment with 20 μM HC3 reduced incorporation into PC in uninfected cells, whereas 200 μM treatment resulted in a dramatic reduction in choline incorporation into PC independent of HCV infection (Figure 4A), validating the inhibition of choline uptake. While the transcript expression of CTL1 was lower with HCV infection after 24 h, treatment with HC3 increased expression to basal levels, with no changes in cells not treated with HC3 (data not shown). Given that we used a 24 h infection model, we sought to determine the effects on early RNA replication. In the presence of 200 μM HC3, the relative amount of HCV was ∼ 2.5 times higher than that of control HCV-infected cells (Figure 4C). Therefore, limiting the availability of choline is associated with an increase in viral replication early during infection in FBS-cultured Huh7.5 cells.

**Figure 4.**
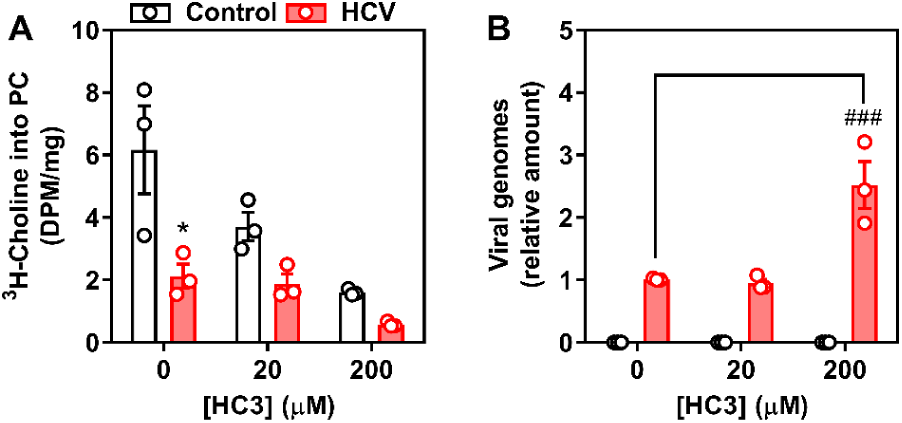
Inhibition of choline uptake augments HCV replication after 24 h. FBS-cultured Huh7.5 cells were infected at an MOI of 2 for 24 h in the presence or absence of 20 or 200 μM hemicholinium-3 (HC3) to inhibit choline uptake. A) [^3^H]-choline incorporation into PC, B) HCV RNA expression as an indication of viral replication (shown relative to infected vehicle control cells) and normalized to the average of *β actin* and *HSP90*. Data are mean±SEM and are represent of 3 independent experiments.

### 3.4. Limiting choline availability during HCV infection of HS-cells inhibits viral infectivity

There is evidence that Huh7.5 cells, when chronically cultured with HS, rather than typical FBS, behave more like human hepatocytes while still in culture. Moreover, the response to HCV infection is also dramatically different. Using cells chronically cultured in HS, we observed no difference in the transcript expression of CTL1 or CTL2 (Figure 5A), as well as no difference in the amount of CTL1 protein (Figure 5B), 4 days post-HCV infection (MOI = 1). To test whether choline availability altered viral replication and infectivity at a later time point, HS-cultured cells were infected with HCV (MOI = 1) and treated with HC3 72 h post-infection. In this culture system, there was no effect of choline uptake inhibition on viral replication (Figure 5C); however, the HCV virus secreted from cells that had been treated with the choline uptake inhibitor were significantly less infectious (Figure 5D).

**Figure 5.**
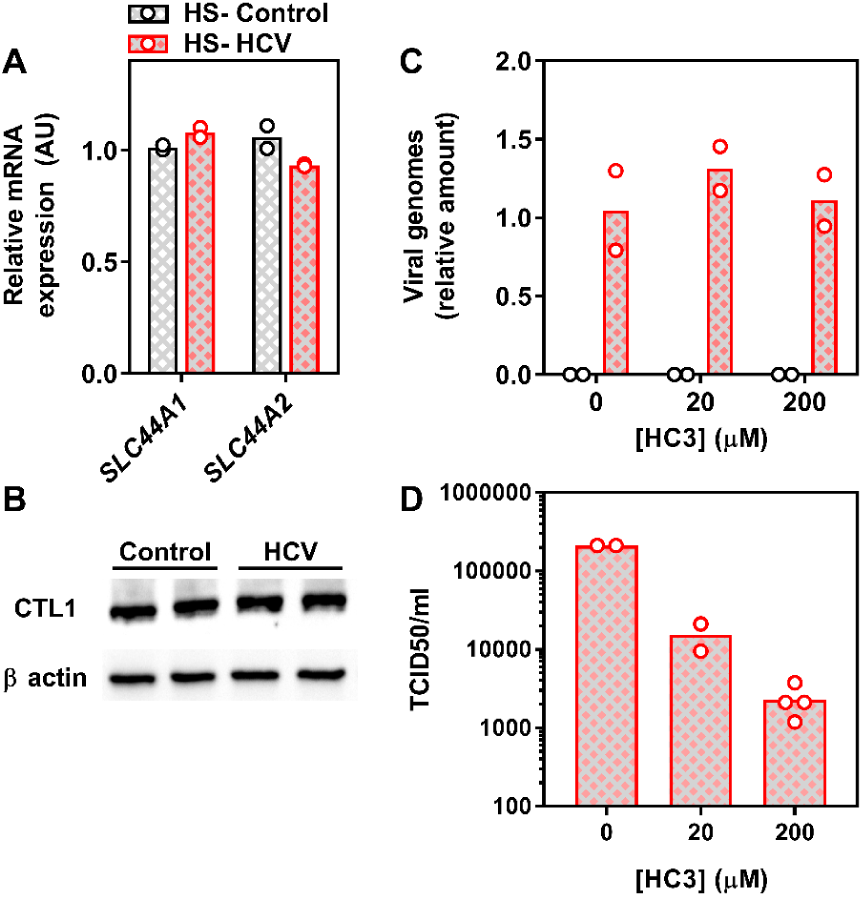
Choline transporters and viral replication is unaltered in human serum-cultured cells. Human serum (HS)-cultured Huh7.5 cells were infected at an MOI of 2 for 96 h and A) choline transporter expression determined, shown relative to non-infected control cells and normalized to the average of *β actin* and *HSP90*. B) CTL1 protein expression. Infected and non-infected cells were cultured in the presence or absence of 20 or 200 μM hemicholinium-3 (HC3) for the final 24 h to inhibit choline uptake before C) intracellular viral RNA was determined, as shown relative to infected vehicle control cells and normalized to the average of *β actin* and *HSP90*. D) TCID50 of secreted virus harvested in the media. Data are mean and are representative of 2 independent experiments performed in duplicate.

## 4 Discussion

Hepatic choline can be used for phospholipid synthesis or can enter the one carbon pathway via its oxidation to betaine. While numerous studies have focused on hepatic choline metabolism, few have looked to address the initial cellular uptake of choline. Here, we provide evidence that in the Huh7.5 human HCC cell line, two choline transporter families are expressed, the low-affinity broad spectrum OCT family and the intermediate affinity CTL family. However, on a functional level, when [^3^H]-choline uptake measurements were fit into a Michaelis-Menten model, it was suggestive of an intermediate affinity for choline (*K*_m_ of 66.8 ± 9.1 µM) thus pointing to the CTL transporters [35]. Moreover, when the contribution of both CTL and OCT transporters was assessed through selective pharmacological inhibition it was clear that only HC3-sensitive (CTL-mediated) transport is present in Huh7.5 cells [34,36]. Interestingly, while OCT family members seemed to play a minor role in choline uptake, the relevance to human hepatocytes *in vivo* warrants further investigation [37,38].

HCV infection leads to widespread hijacking of the host-cell machinery thereby dysregulating a myriad of pathways to allow for its sustained chronic infection of the host [39-43]. Having an innate dependence on enzymes involved in lipid pathways at every stage of its replicative cycle, the link between HCV and lipid metabolism has been well established [24,44,45]. Although entry mechanisms are still not fully understood, it is known that HCV relies on a multitude of receptors, including CD81 and Claudin-1, but also receptors involved in lipoprotein uptake such as LDL-R[46] and SR-BI [47]. The association, although unclear if it is a fusion or a coating with ApoB-100 and ApoE containing lipoproteins (such as VLDLs), is believed to be essential to the formation of highly lipidated infectious lipoviral particles, although a trans-Golgi endosomal pathway has also been proposed [48]. The necessity for VLDL production and how it informs the biogenesis of lipoviral particles was demonstrated by showing that the inhibition of ApoB-100 [49,50] and Apo-E [51,52] in infected cells reduced the number of secreted infectious virions. Our study aimed to address another aspect of lipid metabolism in HCV life cycle, the uptake of choline and its incorporation into PC.

Infection with (+)strand RNA viruses, such as HCV, has been shown to cause an increase in PC amounts at the site of viral replication [16]. Conversely, it was demonstrated that upon HCV infection, the total amount of PC was reduced relative to non-infected cells [53]. The latter is entirely consistent with the lower incorporation of choline into PC we observe 24 h post-infection. Interestingly, HCV requires an interaction with CHKα, the initial enzyme responsible for the phosphorylation of choline in the CDP-choline pathway [53] and there is evidence that CHKα activity, but not its PC-synthesizing function is critical.

The biological significance of lower PC levels in HCV-infected cells remains unclear. *In vivo*, the CDP-choline pathway accounts for approximately 70% of PC, whereas the remaining levels are produced via PEMT and the methylation of PE. Upon HCV infection, there is a reduction in PEMT transcript and protein expression [53]. Our results add to this narrative by demonstrating a reduction in the choline transporter CLT1 after 24 h of HCV infection (Figure 3), which was associated with a decrease in choline uptake and flux through the CDP-choline pathway (Figure 2). Together, this may explain the lower levels of PC. However, our results also provide a snapshot of what is happening at 24 h post-infection. Additionally, it will be important to consider not only the content, but the fatty acid composition of PC species, which also change with HCV infection [54].

Given that catalytic inhibition of CHKα and not necessarily flux through the CDP-choline pathway has been tied to viral replication [53], it was somewhat surprising that the inhibition of choline uptake augmented viral RNA replication after 24 h, albeit slightly. Inhibiting PC synthesis via blockade of the rate limiting enzyme in the CDP-choline pathway, CCTα, was not shown to effect HCV replication [16,21,53]. One potential explanation for this result and discrepancy might be timing. We chose to identify changes to host cell metabolism and viral replication after 24 h, where previous work has only looked at 72 h and beyond. It might also be that limiting choline uptake and availability drives up the expression and activity of other PC-generating pathways (PEMT and PC degradation via phospholipases) to scavenge the PC necessary for viral replication. Interestingly, phospholipase D, which cleaves existing PC to yield free choline and phosphatidic acid, is upregulated in an *in vitro* model of HCV [55].

The human Huh-7 and related cell lines have long been used to address key biological questions with regards to HCV life cycle. While typical culture conditions include the use of standard 10% FBS, it has recently been demonstrated that culturing of Huh7.5 cells in media supplemented with HS, dramatically shifts the cellular and metabolic phenotype of the cells [25,26]. We sought to take advantage of this model system to interrogate the role of HCV infection on the expression of choline transporters, as well as whether inhibition of choline uptake altered viral replication and release. HS-cultured cells, unlike FBS-cultured cells displayed no differences in transporter expression or viral measures. While this cell model may represent a system that is more physiologically similar to fresh primary human hepatocytes, there are potential explanations for the divergent observations. Firstly, the metabolic demands of the cells differ dramatically since FBS-cultured cells divide readily and HS-cultured cells do not. This calls into question whether HCV differentially competes for cellular nutrients such as choline in proliferative vs. quiescent cells. Secondly, we chose to assess early HCV infection (24 h) in our FBS-cultured cells, whereas HS-cultured cells were probed after 4 days of infection. There is the potential that choline transporter expression (CTL1) may be lower initially in HS-cells, but rebounds 96 h post-infection. Similarly, it remains possible that at 96 h post-HCV infection in FBS-cells, CTL1 expression may return to baseline. Lastly, while the 24 h timing of HC3-mediated choline uptake inhibition was identical between FBS- and HS-cells, it is possible that a more prolonged uptake inhibition, such as the entire 96 h infection, may serve to alter HCV replication. Interestingly, in HS-cultured cells, limiting choline availability did not hinder viral replication (Figure 5C), but did significantly inhibit the production of infectious virions (Figure 5D).

In addition to possible timing issues, another consideration is our strict focus on PC metabolism. Choline, once transported into the hepatocyte, can enter into the mitochondria and become oxidized to betaine as the first committed step toward its incorporation as a methyl group donor (for DNA and protein methylation). Along these lines, an initial characterization of hepatic choline metabolic fate has been reported in primary rat vs. a documented HCC cell line, which highlighted the alternative fates of choline toward each arm of the phospholipid and methylation pathways [34]. It will be important for future work to consider the full metabolic consequence of altered choline homeostasis during all stages of *in vitro* HCV infection.

## 5 Conclusions

In conclusion, we demonstrate that short term HCV infection of FBS-cultured Huh7.5 cells results in a substantial impairment of choline uptake and incorporation into PC. This is potentially mediated in part by the lower expression of the main choline transporter CTL1. We also show that when choline uptake is inhibited, this paradoxically slightly, but significantly potentiates viral replication. Finally, in chronically HS-cultured Huh7.5 cells, HCV infection had no effect on choline transporter expression; however, choline uptake inhibition significantly impaired the infectious nature of the virus particles. Taken together, these results demonstrate the potential importance of choline metabolism in HCV and potentially other hepatic viral infections.

## Supplementary Materials

The following are available online at www.mdpi.com/xxx/s1, Table S1: qPCR primer sequences

**Table S1.**
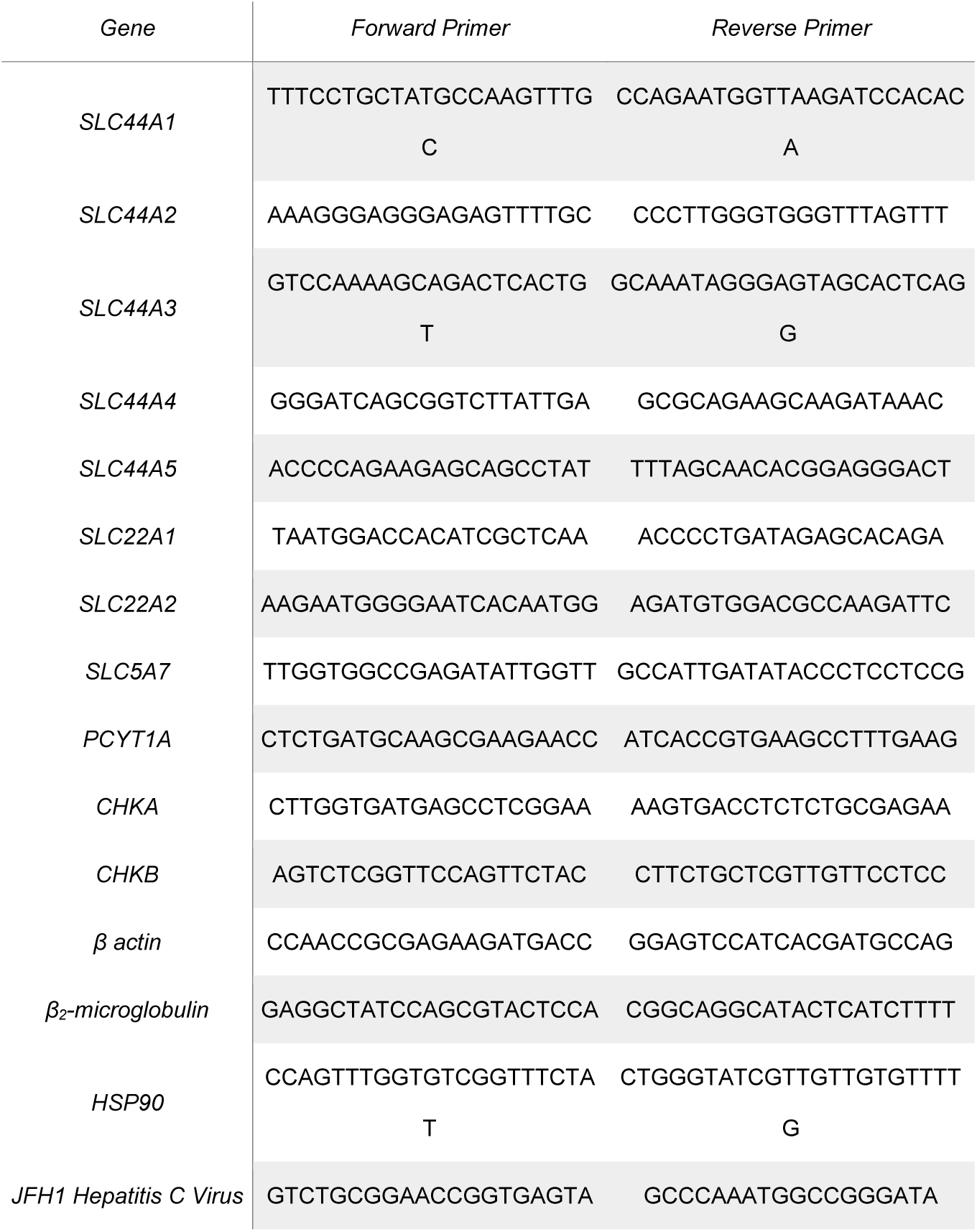
qPCR primer sequences. Tables should be placed in the main text near to the first time they are cited.

## Author Contributions

Conceptualization, Kaelan Gobeil Odai and Morgan D. Fullerton; Formal analysis, Kaelan Gobeil Odai, Rineke Steenbergen and Morgan D. Fullerton; Funding acquisition, Morgan D. Fullerton; Investigation, Kaelan Gobeil Odai, Conor O’Dwyer, Rineke Steenbergen, Tyler A. Shaw, Tyler M. Renner, Peyman Ghorbani, Mojgan Rezaaifar and Shauna Han; Resources, Angela M. Crawley, Rodney S. Russell, John P. Pezacki and D. Lorne Tyrrell; Supervision, Marc-André Langlois, John P. Pezacki, D. Lorne Tyrrell and Morgan D. Fullerton; Writing – original draft, Kaelan Gobeil Odai and Morgan D. Fullerton; Writing – review & editing, Kaelan Gobeil Odai, Conor O’Dwyer, Rineke Steenbergen, Tyler A. Shaw, Tyler M. Renner, Peyman Ghorbani, Mojgan Rezaaifar, Shauna Han, Marc-André Langlois, Angela M. Crawley, Rodney S. Russell, John P. Pezacki, D. Lorne Tyrrell and Morgan D. Fullerton.

## Acknowledgments

We would like to thank Nicholas LeBlond and Rebecca Yaworski for helpful discussions.

